# Trans-acting Determinants of Gene Expression: Effects of Transcription Factor Affinity, Abundance, and Localization

**DOI:** 10.64898/2026.03.10.710832

**Authors:** Maria Lopez-Malo, Sebastian J. Maerkl

## Abstract

Transcription factors (TFs) regulate gene expression by binding cis-regulatory DNA elements, yet how trans-regulatory characteristics such as TF affinity, concentration, and localization interact with cis-regulatory elements remains largely unclear. We systematically analyzed TF affinity mutants across abundance, and localization states and found that promoter binding-site strength most readily modulated expression levels, followed by TF localization and concentration, while affinity variations were mainly buffered. We further uncover performance trade-offs between TF abundance, localization, and affinity. Together, these results reveal how trans and cis factors collectively shape gene-regulatory output.

## Main

Transcription factors (TFs) control gene expression by binding specific DNA sequences, yet the complex interplay between trans-acting properties such as TF affinity, concentration, and localization, and cis-regulatory binding sites remains poorly understood. Cis-regulatory properties have been well studied giving rise to quantitative models of gene regulation [1, 2] and recent work is gaining insights into the functional relevance of low affinity binding site clusters and overlapping binding sites [3, 4, 5]. Trans-properties on the other hand are less well understood [6, 7] despite growing evidence of their importance in central biological processes such as complex regulatory functions, development, and disease [8, 9, 10, 11, 12]. Quantitative studies describing the effect of trans-properties remain rare due to the difficulties associated with precisely modulating and quantifying TF trans-properties *in vivo* [13]. Particularly the contribution of TF affinity has not been investigated because affinity and specificity can’t be easily decoupled. Trans-properties are central to gene regulatory function and unlike changes in cis-regulatory elements, changes in trans-properties will affect the entire regulatory network of a given TF.

To interrogate this complex parameter space we constructed a large combinatorial strain library in *S. cerevisiae* consisting of combinations of eight Zif268 affinity mutants [14] expressed at four different abundance levels using constitutive promoters of different strengths [15], and tested their activation potential on three target promoters with different consensus binding site numbers [3] (Figure 1A). In total, we generated 172 yeast strains to measure the effect of these various trans- and cis-properties. We furthermore modulated TF localization using the *β*-estradiol–inducible Z_3_ EV system [16]. *Z*_3_*EV* is a synthetic TF that consists of the 3 zinc-finger DNA binding domain of the mouse transcription factor, Zif268, a human estrogen receptor (ER) that drives nuclear translocation in the presence of *beta*-estradiol, and a VP16 domain that recruits transcriptional machinery to activate transcription (Supplementary Figure 1 and 2). Zif268 affinity mutants were taken from previous work where we demonstrated that it was possible to tune Zif268 affinity independently of sequence specificity *in vitro*. By mutating amino acid residues that make non-specific contact with the DNA backbone, we obtained new variants covering a range between 2-fold increased and 5-fold reduced affinity without affecting sequence specificity [14]. Gene expression levels were quantified over-time using GFP as a readout. In order to quantitate TF expression levels we generated m-Scarlet - *Z*_3_*EV* fusions. To check if the position of mScarlet had an effect on *Z*_3_*EV* activity we constructed 3 variants of the wt TF-mScarlet fusion: no-tag, N-tag and C-tag, expressed under the 4 constitutive promoters mentioned above. We ultimately chose the C-tag fusion variant as it exhibited the smallest difference compared to the un-tagged TF (Supplementary Figure 3).

**Figure 1.**
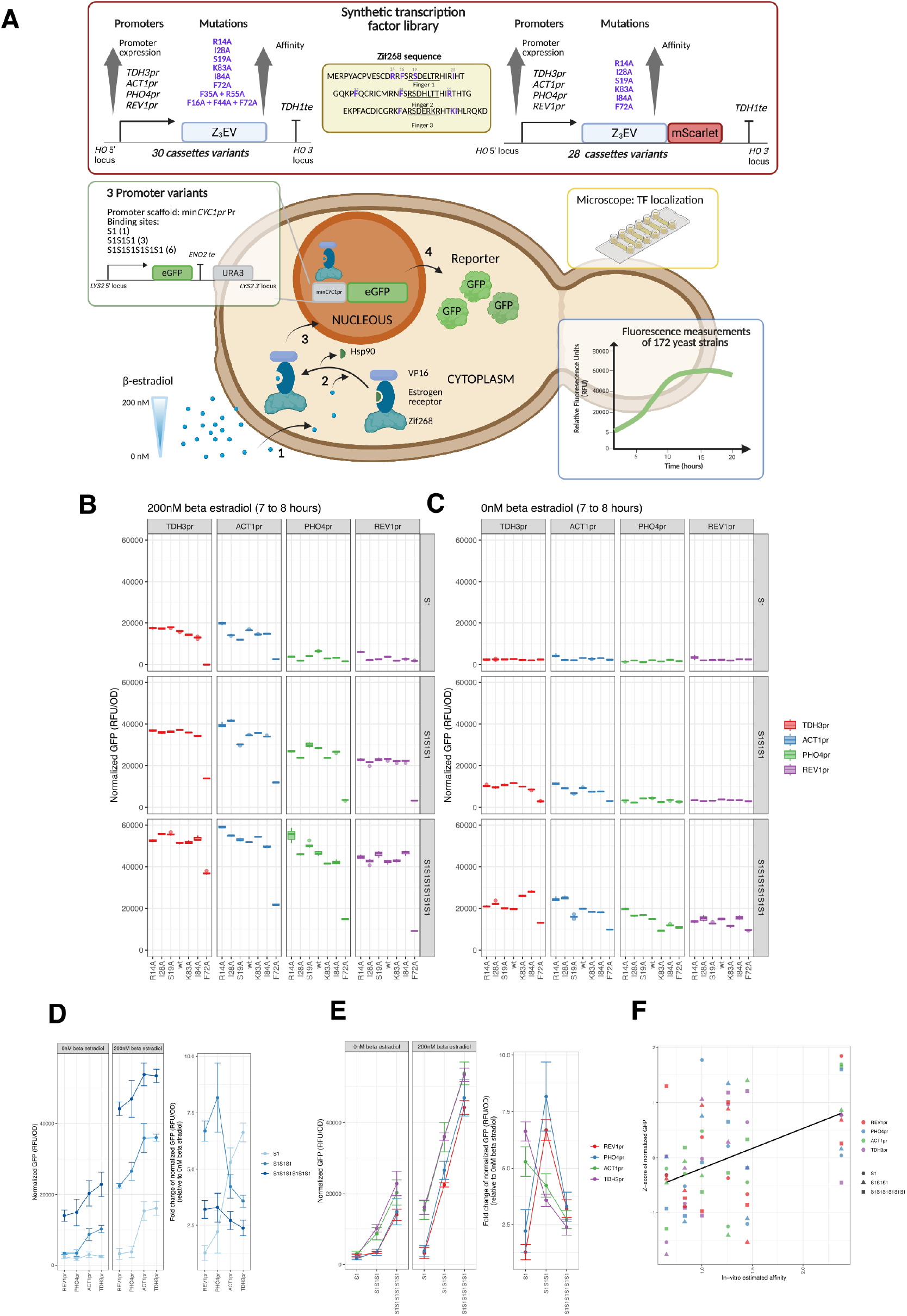
A) Schematic overview of the experimental design including a description of the 172 combinatorial yeast stain library generated in this study. Shown are the various combinations of promoters driving transcription factor mutant expression, the target promoters with 3 different cis-regulatory strengths, and the *beta*-estradiol system for controlling TF localization. All strains are measured over-time on a plate reader (OD and fluorescence) and we used microfluidic devices to quantify TF localization. B) GFP expression results for all Zif268 versions (R14A, I28A, S19A, wt, K83A, I84A and F72A), expressed under 4 different promoters *TDH3pr, ACT1pr, PHO4pr* and *REV1*, binding 1, 3 or 6 binding sites measured between 7 and 8 hours 200nM *beta*-estradiol induction and C) without induction with *beta*-estradiol. Box plot of normalized GFP expression (n=9, central line = median, box = interquartile range (Q1–Q3), whiskers = data range (up to 1.5×IQR), dots = outliers). D) Mean of normalized GFP expression (n=9) measured from 7 to 8 hours after 200nM *beta*-estradiol induction and and without induction with *beta*-estradiol of all Zif268 affinity variants (excluding F72A mutant) binding 1, 3 or 6 binding sites expressed under 4 different promoters *TDH3pr, ACT1pr, PHO4pr* and *REV1*, and the fold change relative to 0nM *beta*-estradiol. E) Mean of normalized GFP expression (n= 9) measured from 7 to 8 hours after 200nM *beta*-estradiol induction and and without induction with *beta*-estradiol of all Zif268 affinity variants (excluding F72A mutant) expressed under 4 different promoters *TDH3pr, ACT1pr, PHO4pr* and *REV1* binding 1, 3 or 6 binding sites and the fold change relative to 0nM *beta*-estradiol. (D-E: points represent the mean and error bars indicate the standard deviation (SD)). F) GFP fluorescence values (excluding F72A mutant) were normalized using z-score transformation to allow comparison between *in vivo* and *in vitro* estimated affinity [14]. Linear regression fit (*y* = 1.26 + 0.276*x, R*^2^ = 0.199, *p −* value = 8.4 × 10^*−*5^).

**Figure 2.**
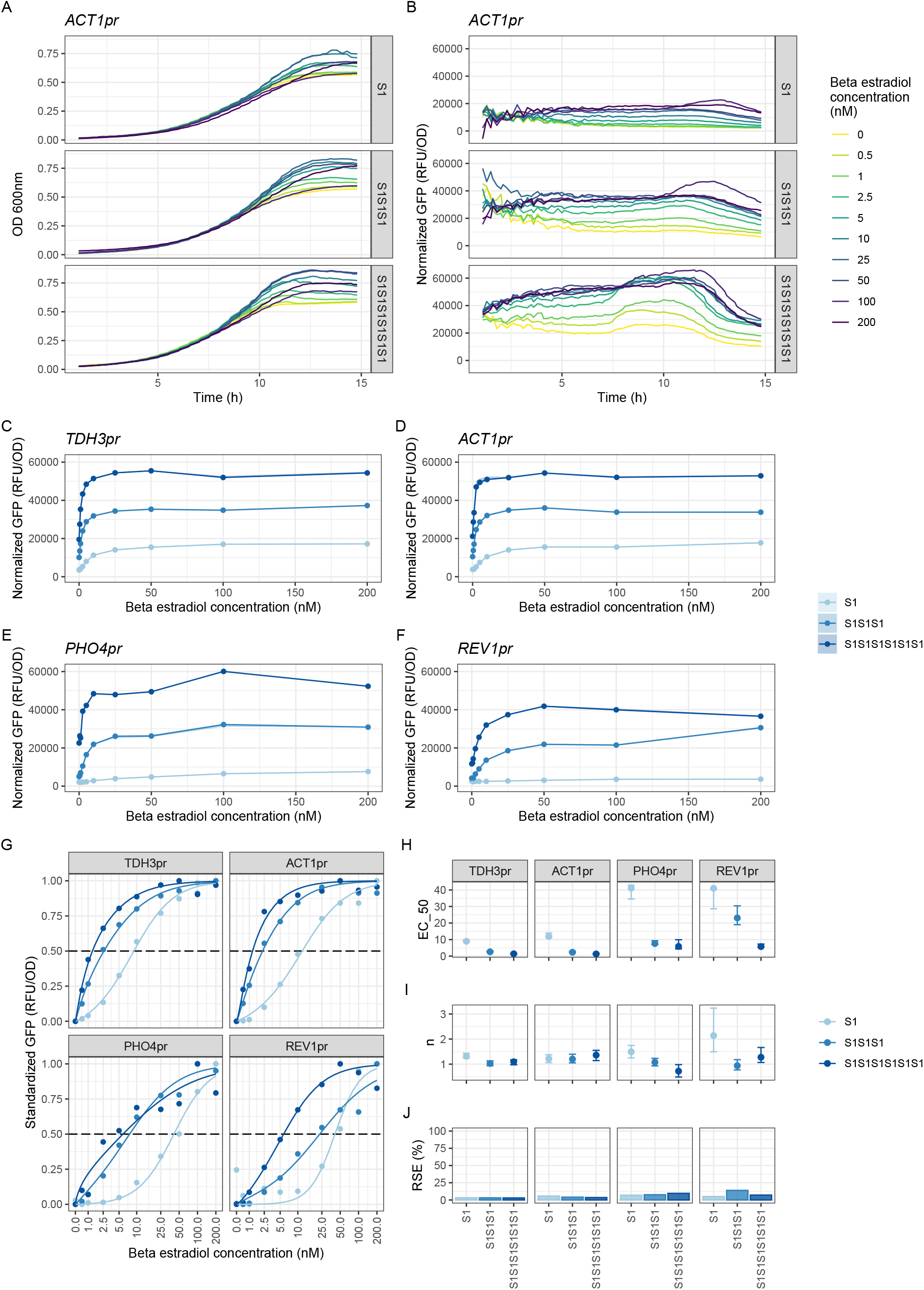
Modulating nuclear localization by *beta*-estradiol titration. Representative examples of A) the OD600 and B) the normalized GFP (RFU/OD600) time courses of wt Zif268 expressed by the *ACT1pr*, under 10 different concentrations of beta estradiol. Lines represent the mean of each *beta*-estradiol concentration (n=9). C-F) Dose-response curves of the normalized GFP values measured between 7 and 8 hours after *beta*-estradiol induction, of the wt version of Zif268 expressed under the C) *TDH3*, D) *ACT1pr*, E) *PHO4pr* and F) *REV1* promoters, binding 1, 3 or 6 binding sites, under 10 different concentrations of *beta*-estradiol. Points represent the mean (n=9). G) The same data as in C-F normalized, plotted on a log axis and fit with a hill function. H) Hill function fit parameters: *beta*-estradiol concentration required for half-maximal response (EC_5_0), the Hill coefficient (n), and the Residual standard error (RSE).

We first assessed the impact of TF affinity and abundance in the context of different cisregulatory binding site strengths. To do so we measured reporter gene expression under fully nuclear localizing (200nM *beta*-estradiol) (Figure 1B, Supplementary Figure 4A) and cytoplasmic conditions (0nM *beta*-estradiol) (Figure 1C, Supplementary Figure 4B). We quantified GFP levels during exponential growth normalized by cell-density (OD) and averaged the resulting values over a 1 hour window. The resulting measurements were robust across technical repeats as well as different time-windows (Supplementary Figure 5).

TF abundance has been known to be an important parameter in development and disease processes, and recent work identified genes that appear to be either sensitive or robust to TF dosage, although the mechanistic details are not yet understood. We modulated TF abundance by expressing our TF variants under 4 constitutive promoters covering a wide range of expression levels ranging from very low expression (REV1pr) to very high expression (TDH3pr) and tested these TF abundances for their ability to activate target promoters of different strengths. We validated that TF abundances indeed correspond to the expected expression levels, by quantifying mScarlet-TF fusions, although it was not possible to detect the difference in expression levels of the two lowest promoters (REV1pr and PHO4pr) as the resulting TF levels were below the detection threshold (Supplementary Figure 6).

We found that all target promoters exhibit TF dosage dependence, but only the weakest cis promoter was particularly sensitive to abundance levels, showing step-function like behavior in that it was not activated by the two lowest TF abundance levels, but was activated by the two higher abundances (Figure 1D). Output of the medium (3S1) and strong (6S1) cis target promoters could be tuned by TF abundance as long as abundance levels remained below the saturating threshold of the two highest TF abundance levels (ACT1pr and TDH3pr) which saturated expression levels for all cis target promoter strengths. The highest fold changes were achieved by the weakest cis target promoter (1S1) in combination with the highest TF abundance, as well as the intermediate strength cis target promoter (3S1) driven by the lowest TF abundance levels. This shows that two completely different strategies can be used to achieve high-fold changes: weak cis promoters coupled with high TF abundance levels or strong cis promoters coupled with low TF abundance levels. Our results also show that cis promoter strength and TF abundance must be carefully balanced. The strongest cis target promoter (6S1) exhibited the lowest fold-change results. In all cases fold-change was limited by TF leakage into the nucleus which is discussed in more detail below.

Standard cis-modulation of target promoter strength achieved by far the widest dynamic range, particularly in combination with low TF abundance levels (REV1pr and PHO4pr) (Figure 1E). Our weakest cis target promoter (1xS1) is still relative strong in that it contains one consensus binding site, it should therefore be possible to access an even higher dynamic range in combination with high TF abundance levels by further reducing cis target promoter strength under these conditions. One significant drawback of strong cis promoters, particularly in combination with high TF abundance is that promoter leakage sets in. We observed two sources for promoter leakiness in the 0nM *beta*-estradiol condition. First the strong target promoter (6xS1) became leaky, likely due to non-specific binding and activation via other TFs as the output was significantly above baseline for all TF abundance levels. The combination REV1pr - F72A in this condition can be considered to be close to a negative control due to the limited functionality of the F72A mutant and its low abundance. We observed this type of leakiness in our previous work as well [3].

In this dataset we were able to identify a second source of leakiness which is due to TFs leaking into the nucleus under the non nuclear-localizing condition (0nM *beta*-estradiol). This occurred in fact at all TF abundance levels tested and correlated with TF abundance (higher abundance leading to higher leakage). This effect was most apparent in the strong 6xS1 target promoter which was sensitive to even small TF leaks. Induction was still apparent in the medium strength target promoter 3xS1 but only for the two highest TF abundance levels, and aberrant induction, despite TF leaking into the nucleus, was completely eliminated with the low strength 1xS1 target promoter.

Although our TF affinity variants cover a 4 fold-range in affinities we observed only a slight relationship between TF affinity and gene expression levels under any of the TF abundance and target promoter strength combinations that were tested (Figure 1F). The only exception was the lowest affinity variant (F72A) which failed to activate the single target site promoter (1xS1) under all TF abundance levels, but was able to activate promoters with medium (3xS1) and high (6xS1) number of binding sites in an abundance dependent manner (Figure 1B). This indicates that WT Zif268 is overall well buffered against affinity mutations in that affinity changes did not result in corresponding changes in gene expression.

This is somewhat surprising, as a priori we expected a stronger effect of TF affinity on activation levels, similar to what was observed for the lowest affinity mutant. Nonetheless, it is possible to tune gene expression by modulating TF affinity, but the window in which affinity and activity correlate appears to be narrow, as further reduction in affinity through double mutants completely ablated function (Supplementary Figure 4). It is important to note that we focused on mutations that solely affect affinity but not specificity, which is considerably different from mutations that would also affect specificity. For example, adding another Zn-finger DNA binding domain to Zif268 would lead to a large increase in DNA binding affinity but would also change the DNA binding specificity of the TF [17]. The observed buffering could be explained by a complex interplay of affinity to specific as well as non-specific DNA. For example, an affinity increasing mutation increases the affinity to the specific target sites which should give rise to increased gene expression levels. But this might be counterbalanced through a concurrent increase in affinity to non-specific DNA which could result in a TF titration effect, reducing the effective concentration of the TF [18].

TF localization is a second stage of modulating and regulating effective TF abundance by controlling the number of TFs present in the nucleus. We therefore assessed how precisely gene expression could be regulated via TF localization by performing a titration of *beta*-estradiol controlling nuclear localization of the WT Zif268 TF (Figure 1A-F,Supplementary Figure 7)). Saturation was achieved at 50 nM *beta*-estradiol for very low TF abundance levels. Higher transcription factor abundance shifted the saturation to lower *beta*-estradiol concentrations (10 nM) and steeper response curves.

To precisely quantitate the dependence on *beta*-estradiol we fit hill equations to the normalized dose response curves (Figure 1G and H). All curves produced a hill coefficient of around 1 except for the REV1pr-1xS1 combination, which is likely due to minimal activation and poor fit. On the other hand, the *beta*-estradiol concentration at which 50% output is reached (EC_5_0) was both a function of TF abundance and promoter strength with higher TF abundance or higher promoter strengths shifting the EC_5_0 to lower *beta*-estradiol concentrations making it more difficult to precisely control gene expression levels under these conditions. It is therefore possible to achieve either a finely controllable dose response (high EC_5_0 values, low TF abundance or low target promoter strength) or more “sensitive” step-like response (low EC_5_0 values, high TF abundance and high target promoter strength). One tradeoff in achieving finely-controllable dose responses with higher EC_5_0 values is that the maximum output and therefore dynamic range tends to be lower than what can be achieved with response curves with lower EC_5_0 values.

Our results show that DNA binding site strength had the largest effect on gene expression levels, followed by TF concentration (Supplementary Figure 8). TF affinity had negligible impact on modulating gene expression, with the exception of a drastically affinity reduced mutant. Regulating gene expression by modulating TF localization was overall robust, enabling high to medium levels of tunability which depended primarily on TF abundance. All of these parameters interacted with one another leading to tradeoffs in either maximum expression levels, dynamic range, and corresponding promoter leakiness and tunability. The overall best performing combination of factors in terms of dynamic range and tunability was achieved with low TF abundance levels driven by the PHO4pr or REV1pr. It is maybe not surprising that Pho4pr performed well, as it defines the expression levels of the endogenous yeast TF Pho4p, but it also suggests that native gene regulatory networks may have evolved towards these optimal parameters. Together, our results reveal how TF abundance, affinity, localization, and target binding-site multiplicity shape the functional landscape of gene regulation and provide a comprehensive framework for engineering transcriptional circuits and provide insights for deciphering the function and evolution of natural gene regulatory networks.

## Material and Methods

### Media and growth conditions

For yeast transformation, cells were incubated at 30°C, with shaking at 250 rpm in yeast extract peptone dextrose (YPD) medium, (Y1375-1KG, Sigma-Aldrich) (10 g/L yeast extract, 20 g/L peptone and 20 g/L glucose). Plates contain 20 g/L of agar (05039-500G, Sigma-Aldrich). All reagents used in this study are listed in Table 1.

Synthetic transcription factor library yeast strains were culture in synthetic complete (SC) medium lacking uracil. 1.92 g of yeast synthetic dropout medium supplement (without uracil) (Y1501-20G, Sigma-Aldrich), and 6.7 g Yeast Nitrogen Base (YNB) without amino acids (Y0626-1KG, Sigma-Aldrich), were dissolved in 960 mL of deionized water and autoclaved. After sterilization we added 40 mL 50% filter sterilized glucose solution to a final volume of 1 L. To cure the transformed colonies, 5-Fluoroorotic Acid supplemented synthetic complete medium (SC 5-FOA) plates were used. To prepare the SC 5-FOA plates, we added 1.92 g yeast synthetic dropout medium supplement (without uracil) (Y1501-20G), 6.7 g Yeast Nitrogen Base without amino acids (Y0626-1KG), and 20 g Agar in deionized water to 880 mL and after autoclaving, we added 40 mL 50% filter sterilized glucose solution, 66 mL 760 *µg/mL* uracil solution, 10 mL 10% 5-FOA solution (EZSolutionTM, 00386260) to a final volume of 1 L.

### Synthetic transcription factor *Z*_3_*EV* plasmid library construction

Transcription factor (TF) variants were ordered as DNA fragments from TWIST Bioscience. The synthetic transcription factor *Z*_3_*EV* library integration cassette consisted of two *HO* homology arms (each 500 bp), promoter variants, *TDH3pr* (680 bp), *ACT1pr* (661 bp), *PHO4pr* (441 bp), *REV1pr* (700 bp), the mouse Zif268 transcription factor variants (270 bp), the ligand binding domain of the human estrogen receptor (885 bp), VP16 activation domain (201 bp), and *TDH1* terminator sequence (224 bp). Alternative versions of all integration cassettes were created adding the yeast codon optimized mScarlet-i (yomScarlet-i) fused to the C-terminus (Figure 1). In addition, for all wt Zif268 integration cassettes N-tagged mScarlet-i versions were constructed.

All integration cassettes were constructed hierarchically by using modular part plasmids following the YeastToolkit (YTK) format [15]. Briefly, part plasmids were generated by integrating DNA fragments via BsmBI goldengate assembly into pYTK001, then multiple YTK part plasmids were assembled into cassette plasmids via BsaI goldengate assembly and then multiple cassette plasmids were assembled to the multi-cassette plasmids via BsmBI goldengate assembly. All plasmids are listed in Table 2. Goldengate reactions were prepared as following: adding the volume necessary to have 100 ng of each plasmid, except for the backbone which was added at 25 ng, 2 µL of 10x T4 ligase buffer (NEB), 1 µL of T4 ligase (NEB), 1 µL of BsaI or BsmBI (NEB), and water to final volume of 20 µL. Thermocycler setup: (BsmBI assembly: 45ºC for 2 min/BsaI assembly: 42ºC for 2 min, 16ºC for 5 min) x 25 cycles, followed by a final digestion step at 60ºC for 10 min and a heat inactivation at 80ºC for 20 min. To assemble the *Z*_3_*EV* no-tag library cassette plasmids (Wildtype TF (wt): pML056, pML080, pML086 and pML092; single mutant TFs: from pML114 to pML131 and from pML193 to pML198; double and triple mutant TF from pML219 to pML222) assembly connector ConLS part plasmid (pYTK002), promoters variant part plasmids or PCR product (pML077, pYTK009, *PHO4pr* PCR product, oligonucleotides are listed in Table 3, and pYTK027), Zif268 transcription factor variants and estrogen receptor (Z3E) part plasmid (from pML058 to pML063, pML066 and pML067), VP16 part plasmid (pSC012), *TDH1* terminator part plasmid (pYTK056), assembly connector ConRE part plasmid (pYTK072) were assembled into the backbone plasmid (pYTK095) via a BsaI goldengate assembly. Z3EV no tag library integration plasmids (wt: pML057, pML081, pML087 and pML093; single mutants: from pML132 to pML149 and from pML199 to pML204; double and triple mutants from pML223 to pML226) were constructed by BsmBI assembly of cassettes plasmids (wt: pML056, pML080, pML086 and pML092; single mutants: from pML114 to pML131 and from pML193 to pML198; double and triple mutants from pML219 to pML222) into the backbone plasmid that contains two *HO* homology arms (pSC017). To assemble *Z*_3_*EV* C-tag library cassette plasmids (wt: pML183, pML184, pML156 and pML217; mutants: from pML150 to pML155, from pML165 to pML170, from pML177 to pML182 and from pML205 to pML210), two PCR products were assembled into the backbone plasmid (pYTK095) via a BsaI goldengate assembly. To amplify the N terminus of the cassette, no tag library cassette plasmids were used as a template to obtain a PCR product that contains the assembly connector ConLS and Zif268 transcription factor variants. To amplify the C terminus from a C-tagged mScarlet version, expressed under *ACT1*, cassette plasmid pML084 (a C-tag plasmid assembled as explained for no tag cassette plasmids but using alternative part plasmids for VP16, pML073, for mScarlet, pML079 and for *TDH1* terminator, pYTK066) was used as a template. This PCR product contains the ER ligand binding domain, VP16 activator, the yeast codon optimized mScarlet-i and the assembly connector ConRE. *Z*_3_*EV* C-tag library integration plasmids (wt: pML191, pML192, pML097 and pML218; mutants: from pML159 to pML164, from pML171 to pML176, from 144 to pML149 and from pML211 to pML216) were constructed by BsmBI assembly of C-tag cassettes plasmids (wt: pML183, pML184, pML156 and pML217; mutants: from pML150 to pML155, from pML165 to oML170, from oML177 to pML182 and from pML205 to pML210) into the backbone plasmid that contains two *HO* homology arms (pSC017). We constructed the *Z*_3_*EV* N-tag cassette plasmids versions only for the wt of Zif268 (pML071, pML082, pML088 and pML094). To do that, the assembly connector ConLS part plasmid (pYTK002), promoters variant part plasmids or PCR product (pML077, pYTK009, *PHO4pr* PCR product and pYTK027), the yeast codon optimized mScarlet-i (pML078), the wt Zif268 transcription factor and estrogen receptor (Z3E) part plasmid (pML070), VP16 part plasmid (pSC012), *TDH1* terminator part plasmid (pYTK056), assembly connector ConRE part plasmid(pYTK072) were assembled into the backbone plasmid (pYTK095) via a BsaI goldengate assembly. *Z*_3_*EV* N-tag library integration plasmids (pML072, pML083, pML089 and pML095) were constructed by BsmBI assembly of cassettes plasmids (pML071, pML082, pML088 and pML094) into the backbone plasmid that contains two *HO* homology arms (pSC017).

For all plasmid cloning we used NEB 10-beta Competent *Escherichia coli* (High Efficiency) (New England Biolabs, Cat #C3019H). We followed the transformation protocol from the manufacturer. We conducted bacterial selection and growth on Lysogeny Broth (LB) plates or in LB medium at 37°C, supplemented with appropriate antibiotic, (chloramphenicol 34 *µg/mL*, ampicillin 100 *µg/mL* or kanamycin 50 *µg/mL*). All plasmids were sequence-verified by Sanger sequencing (Microsynth Ecoli NightSeq).

### Yeast strain construction

For transcription factor (TF) yeast integration, we used CRISPR/Cas9 assisted markerless integration by homologous recombination. Integrating the transcription factor we obtained 64 strains (sML106, sML110, sML118, sML119, sML121, sML122, from sML124 to sML126, from sML154 to sML165,, from sML203 to sML208, from sML230 to sML249, from sML310 to sML322 and from sML362 to sML365), (Table 4). These strains were used as parental strains to subsequently integrate the minimal *CYC1* promoter in the yeast genome. The *HO* sgRNA plasmid (pMC120), CRISPR/Cas9 plasmid (pWS158), and integration cassettes were sequence-verified and digested by EcoRV, BsmBI, and NotI for yeast integration, respectively. The digestions were set up as described Yip *et al* [19], with the difference that we digested with NotI 2 *µg* of the integration cassettes. Alternatively, C-tag integration cassettes were amplified by PCR. After DNA purification (ZYMO DNA clean and concentrator-25 kit), they were transformed and integrated into the parental strain (BY4741) by homologous recombination using the lithium acetate/polyethylene glycol (PEG) method [20]. Briefly, the parental strain BY4741 was grown overnight in YPD medium at 30ºC and shaking at 250 rpm. After overnight culture, the yeast cells for each transformation were diluted to optical density (OD) of 0.175 in 5 mL fresh YPD medium. We prepared 54 *µL* of DNA mixture solution for each transformation, including 2 *µg* integration cassette, 100 ng Ca9/URA3 cassette, and 200 ng sgRNA cassette. The sgRNA coding sequence cassette and Cas9-*URA3* cassette after transformation, were combined to form a plasmid and expressed the Cas9-sgRNA complex to target the integration loci. We conducted the transformant selection on synthetic complete uracil dropout (SC-URA) agar plates and cured the selected colonies on 5-FOA supplemented synthetic complete (SC 5-FOA) plates. We screened transformants for correct integration by colony PCR and verified the PCR product by Sanger sequencing (Microsynth).

For promoter yeast integration we used 3 variants of minimal *CYC1* promoter integration cassettes, pAS019, pML041 and pML042, described in Shahein *et al*.[3]. Briefly, these cassettes consisted of two *LYS2* homology arms (each 500bp), *CYC1* minimal promoter variants (pAS019 one binding site, S1, pML041 three binding sites, S1S1S1 and pML042 six binding sites, S1S1S1S1S1S1), yoEGFP, *ENO2* terminator and *URA3* auxotrophic marker. The three integration cassettes plasmids were digested with NotI and after that the DNA was purified (ZYMO DNA clean and concentrator-25 kit). All promoter library yeast strains were made by homologous recombination of the integration cassettes, into *LYS2* locus, using the lithium acetate/ PEG method. We used 2 *µg* of these 3 integration cassettes to transform the TF genome integrated strains (64 strains), to obtain 184 new strains (From sML107 to sML109, from sML112 to sML114, from sML127 to sML132, from sML136 to sML141, from sML145 to sML153, from sML166 to sML201, from sML212 to sML229, from sML250 to sML309, from sML324 to sML361 and from sML366 to sML368). The transformants were selected in SC medium lacking uracil plates at 30ºC. We screened transformants for correct integration by colony PCR and followed by Sanger sequencing of the colony PCR product (Microsynth).

### Plate reader characterization

The library strains were grown overnight in 3 ml in SC medium lacking uracil at 30ºC shaking at 250 rpm. Cultures were diluted at OD of 0.175 in fresh SC medium lacking uracil. Cells were grown until log phase, OD ≈ 0.8, then diluted to a starting OD of 0.1 – 0.2 in SC medium lacking uracil and SC medium lacking uracil with *beta*-estradiol (Sigma-Aldrich) at 200 nM. Cells were culture in 96-well plates with clear flat bottom (Nunc) and covered with a permeable membrane (Breathe-Easy membrane, Sigma-Aldrich). OD600, GFP and mScarlet were measured every 10 min for 20 – 24 h on a plate reader (BioTek SynergyMx). 96-well plates were incubated at 30ºC with agitation. Media background was subtracted, then the data was normalized by dividing by the OD at each time point. For each strain we preformed 3 independent biological experiments with at least 3 technical replicates.

A *β*-estradiol titration was preformed using the wt Zif268 mScarlet C-tag expressed under the 4 different promoters (From sML289 to sML291 strains, *TDH3pr*, from sML286 to sML288 strains, *ACT1pr*, from sML151 to sML153 strains *PHO4pr* and from sML359 to sML360 strains, *REV1pr*). 10 different *β*-estradiol concentrations were tested, 0 nM, 0.5 nM, 1 nM, 2.5 nM, 5 nM, 10 nM, 25 nM, 50 nM, 100 nM and 200 nM. Yeast cultures and plate reader measurements were preformed as described above. After the *β*-estradiol titration analysis 5 concentrations (0 nM, 1 nM, 5 nM, 50 nM and 200 nM) were chosen to do a deep characterization of Zif268 mutants expressed under *ACT1* (from sML212 to sML229 strains) and *REV1* promoters (from sML341 to sML358 strains).

### Fluorescence microscope analysis

Wt Zif268 expressed under *TDH3pr* and *ACT1pr* containing one binding site (S1) *CYC1* minimal promoter, sML286 and sML289 strains, respectively, were used for transcription factor localization. Yeasts were grown overnight in 3 ml in SC medium lacking uracil at 30ºC and shaking at 250 rpm. Cultures were diluted at OD of 0.175 in fresh SC medium lacking uracil. Cells were grown until log phase, OD ≈ 0.8, then 1 mL of the culture was centrifuged. The pellet was re-suspended with 50 *µL* of fresh media, then 30 *µL* of yeast suspension was loaded into one channel of *µ*-Slide Vi 0.4 (Ibidi, Germany). Then the image before *β*-estradiol induction was recorded. Media was exchanged by aspirating both reservoirs, washing twice with 120 *µL* of fresh SC medium lacking uracil and finally adding 120 *µL* of fresh SC medium lacking uracil with 200 nM *β*-estradiol. The fresh media was slowly filled into one reservoir. After induction the samples were imaged at 10 min, 20 min and 30 min, using a Nikon Eclipse Ti microscope with a 100x oil objective (Nikon type F, Nikon, Japan) and using the NIS-Elements viewer software (Nikon Instruments). Cells were imaged in bright-field, 2 ms of exposure time, and fluorescence mode, 4 s of exposure using mCherry laser. Image data was analyzed with Image J software.

## Author Contribution

MLM performed experiments. MLM and SJM designed experiments, analyzed data, and wrote the manuscript.

## Acknowledgements

This work was supported by a Swiss National Science Foundation Sinergia grant (CRSII5-189910).

