## Supplementary Information for "Trans-acting Determinants of Gene Expression: Effects of Transcription Factor Affinity, Abundance, and Localization"

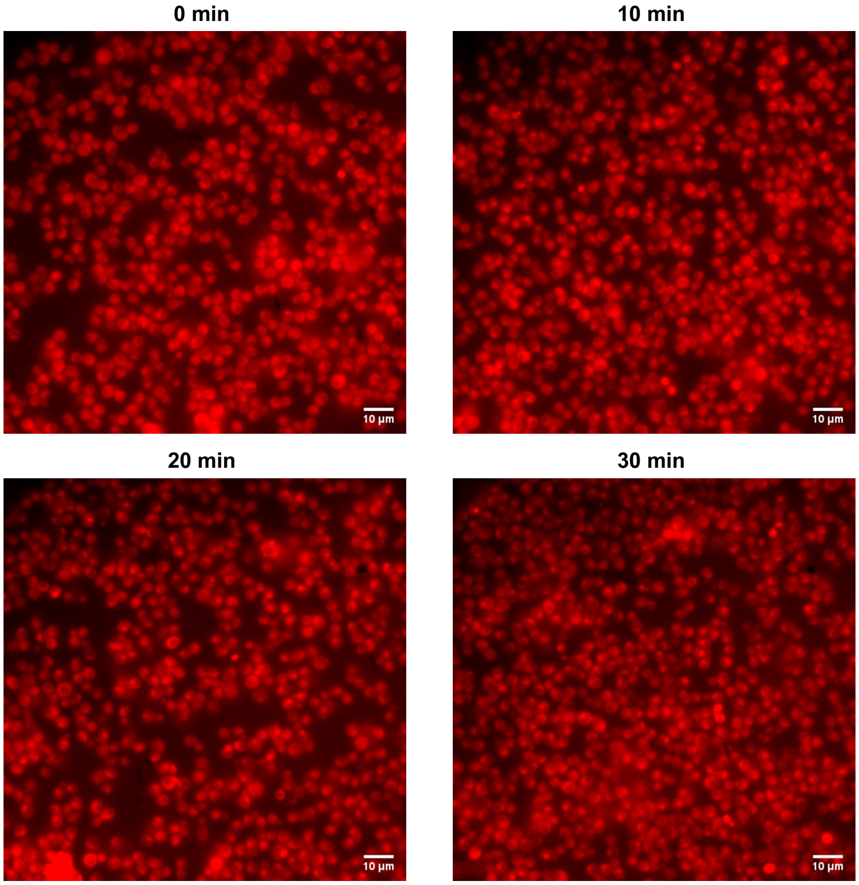


Figure 1: Microscope visualization of $Z_3EV$, expressed under $ACT1pr$, before induction and after 10 min, 20 min and 30 min induction with a saturated concentration of *beta*-estradiol, (200nM).

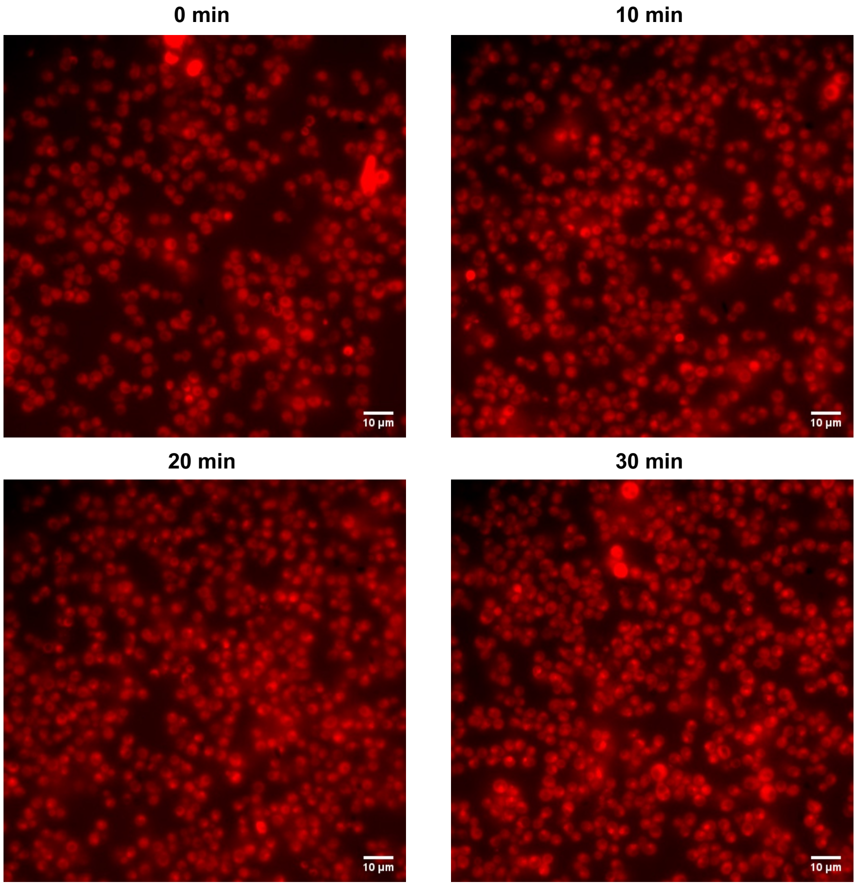


Figure 2: Microscope visualization of $Z_3EV$, expressed under *TDH3pr*, before induction and after 10 min, 20 min and 30 min induction with a saturated concentration of *beta*-estradiol, (200nM).

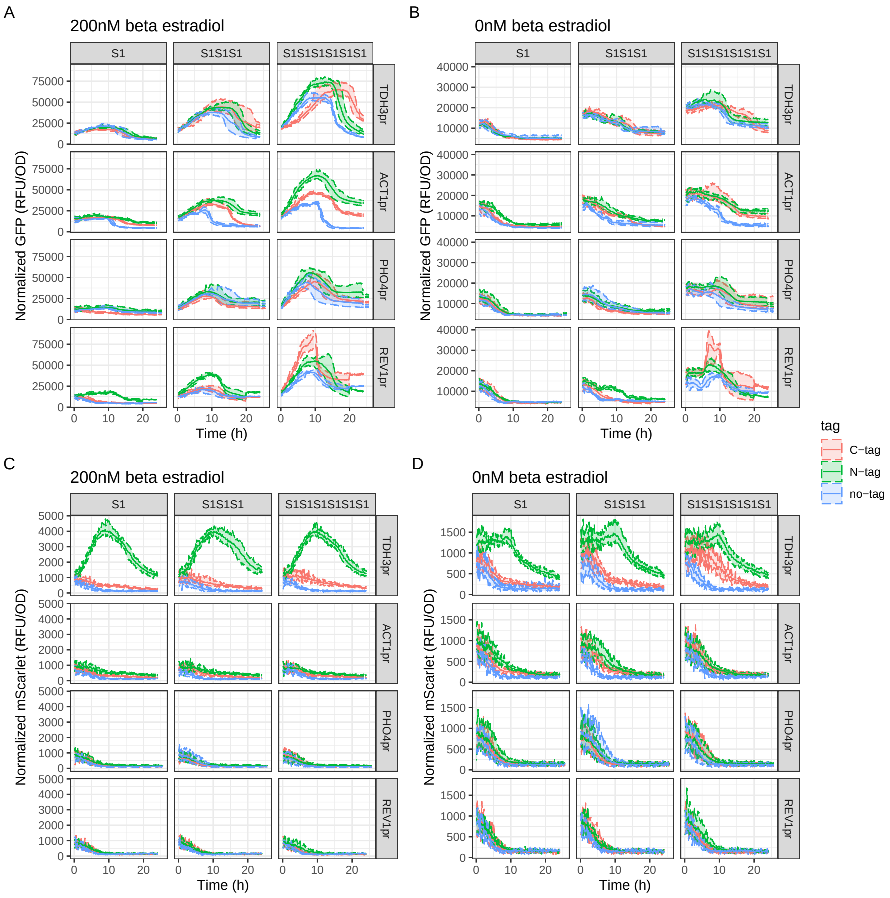


Figure 3: Characterization of C-tag, N-tag and no tag wt $Z_3EV$, expressed under *TDH3pr*,*ACT1pr*, *PHO4pr* and *REV1pr*. GFP expression time courses: A) induced with 200 nM of beta estradiol and B) not induced. mScarlet expression time courses: C) induced with 200 nM of beta estradiol and D) not induced. Dashed lines represent the the maximum and minimum values and solid lines represent the mean (n=9).

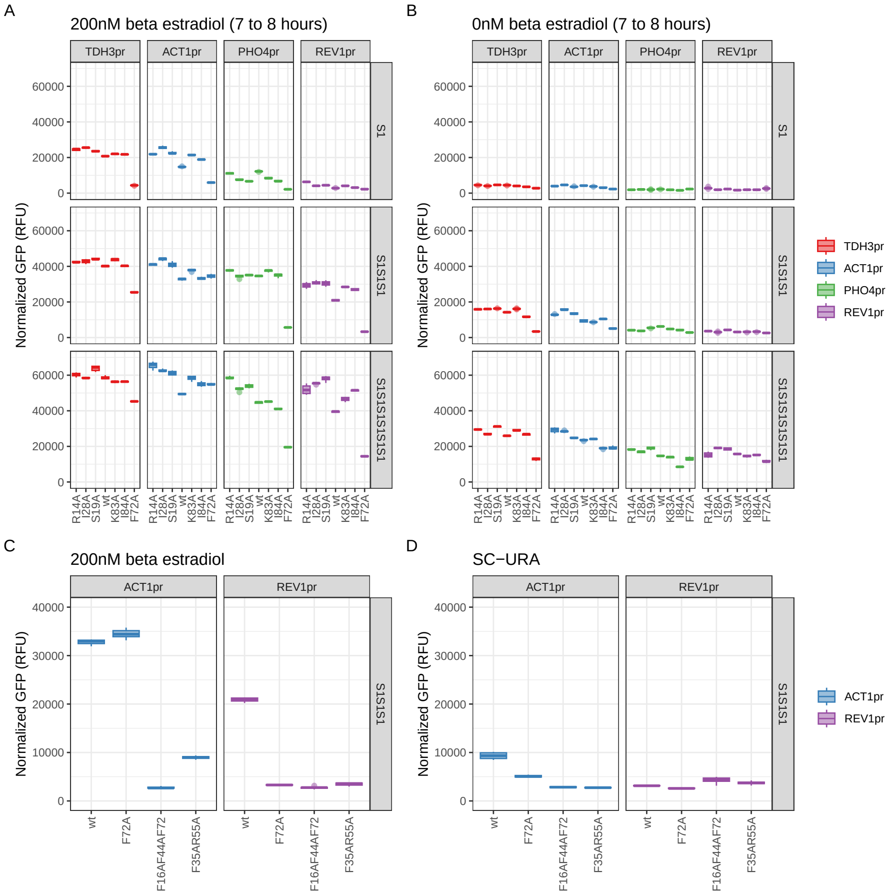


Figure 4: Characterization of no-tagged $Z_3EV$ synthetic transcription library. For all strains, OD600 and GFP were measured every 10 min for 20 – 24 h on a plate reader. Results of the normalized GFP values between 7 and 8 hours for Zif268 versions: R14A, I28A, S19A, wt, K83A, I84A and F72A, expressed under 4 different promoters *TDH3pr*, *ACT1pr*, *PHO4pr* and *REV1*, binding 1, 3 or 6 biding sites, A) after induction with 200nM *beta*-estradiol and B) without induction with *beta*-estradiol. Normalized GFP results between 7 and 8 hours of Zif268 wt version, F72A mutant, double mutant, F35A+R55A, and triple mutant, F16A+F44A+F72A, expressed under *ACT1pr* and *REV1* binding 3 biding sites, C) after induction with 200nM *beta*-estradiol and D) without induction with *beta*-estradiol. Box plot showing the distribution of normalized GFP expression (n=9). The central line represents the median, the box indicates the interquartile range (Q1–Q3), whiskers represent the data range (up to 1.5×IQR), and dots indicate outliers.

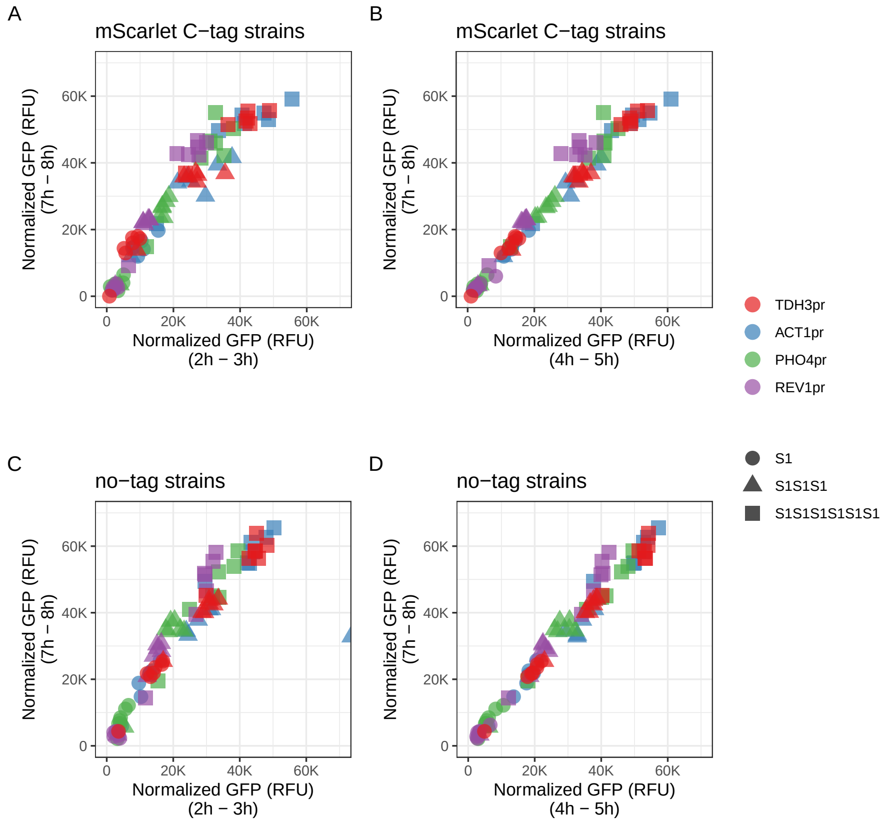


Figure 5: Time correlations of normalized GFP values. A) and B) mScarlet C-tag strains time correlations, and C) and D) no-tag strains time correlations.

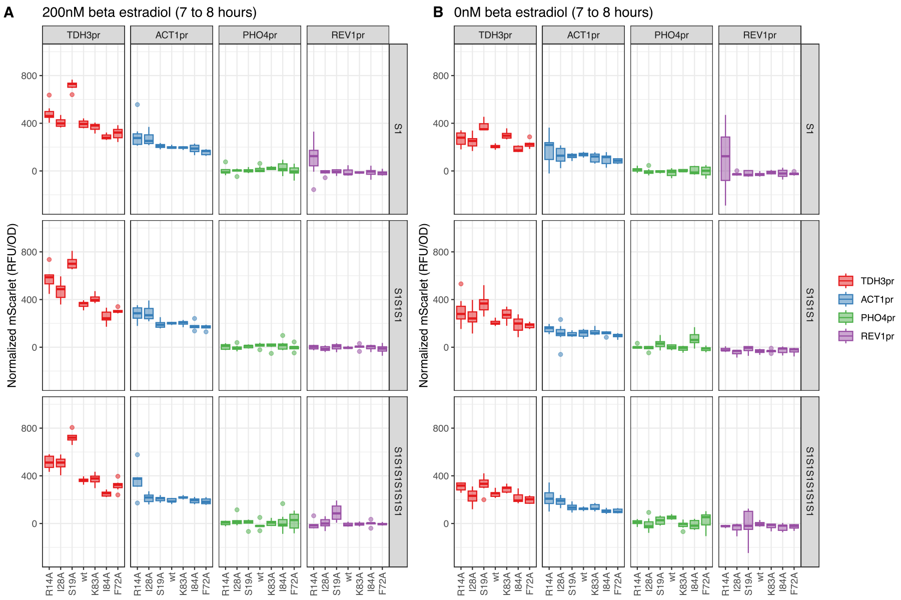


Figure 6: The normalized mScartet expression values between 7 and 8 hours A) after 200nM *beta*-estradiol and B) without *beta*-estradiol induction were plotted as we described for GFP normalized values.

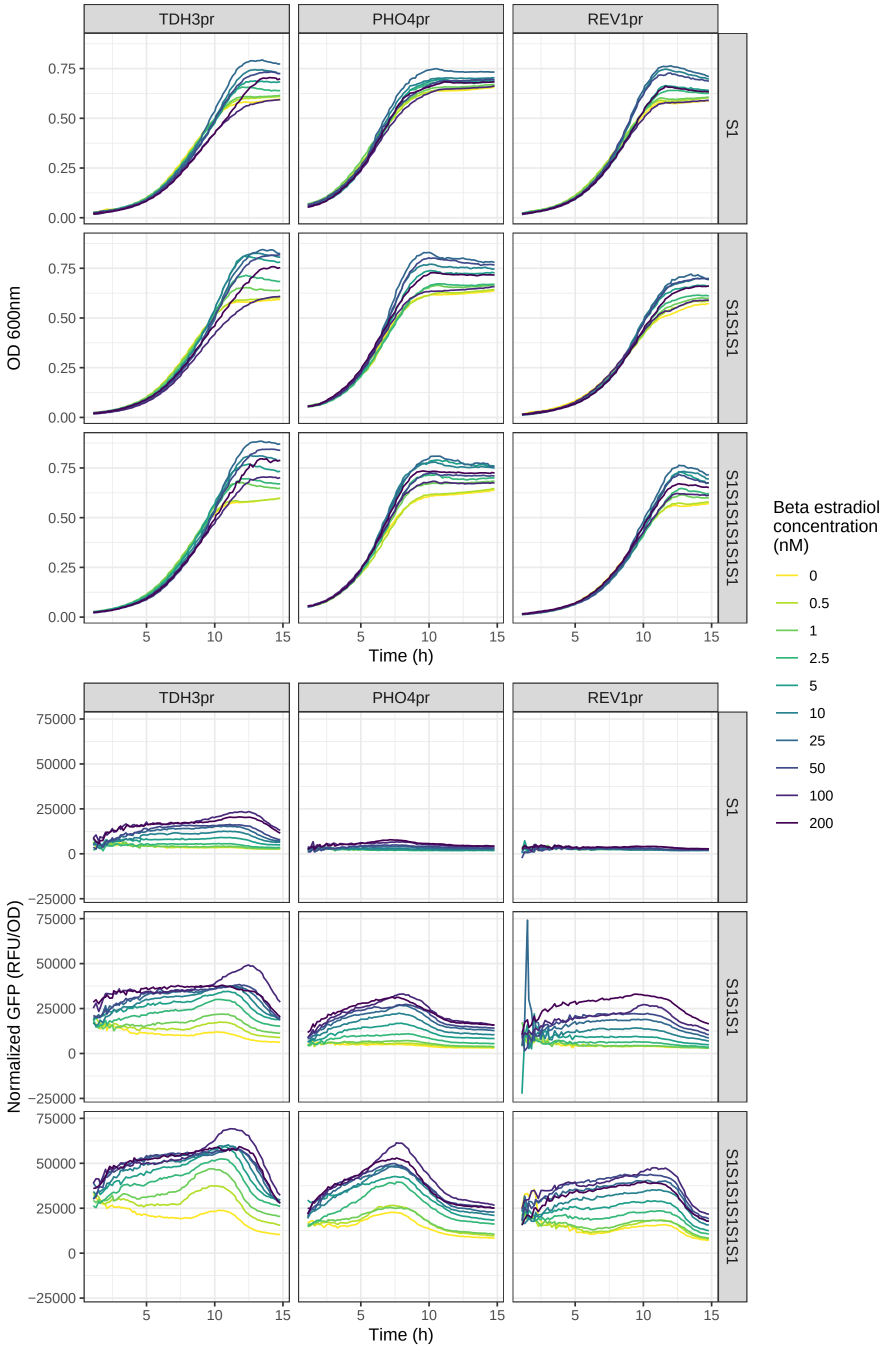


Figure 7: *beta*-estradiol titration results. The OD600 and the normalized GFP (RFU/OD600) time courses of the strains wt Zif268 expressed under *TDH3pr*, *PHO4pr* and *REV1*, under 10 different concentrations of *beta*-estradiol. Lines represent the mean of each *beta*-estradiol concentration.

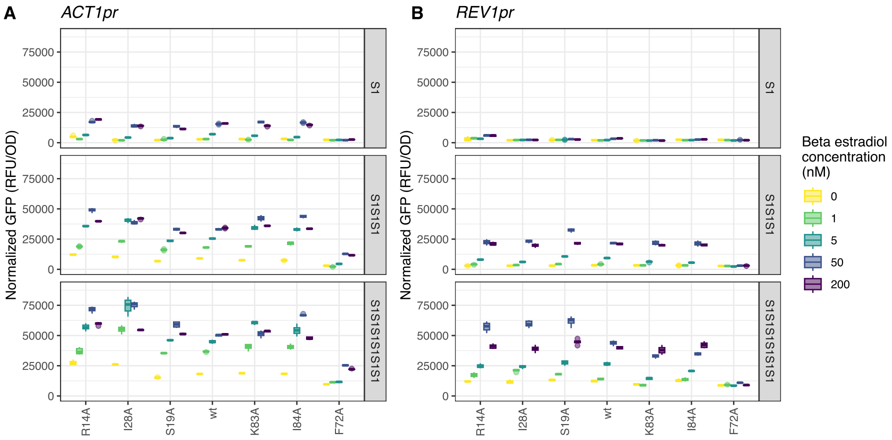


Figure 8: Normalized GFP values of the 7 versions of Zif268, measured between 7 and 8 hours after the addition of 5 different concentrations of *beta*-estradiol, expressed under A) *ACT1pr* and B) *REV1pr*. Box plot showing the distribution of normalized GFP expression (n=9). The central line represents the median, the box indicates the interquartile range (Q1–Q3), whiskers represent the data range (up to 1.5×IQR).
